# Insertional mutagenesis in PLN^R9C^ transgenic mouse: a case report and a review of the literature on cardiac-specific transgenic expression

**DOI:** 10.1101/075671

**Authors:** Alexander Kraev

**Affiliations:** University of Toronto; 27 King's College Circle, Toronto, Ontario, Canada M5S 1A1; Email address

## Abstract

A mouse line with heterozygous transgenic expression of phospholamban carrying a substitution of cysteine for arginine 9 (PLN^R9C^) under the control of α-myosin heavy chain (αMHC) promoter features dilated cardiomyopathy, heart failure and premature death. In this line the transgenic array of 13 PLN^R9C^ expression cassettes, arranged in a head-to-tail tandem orientation, has integrated into the homologous genomic site, the bi-directional promoter of the αMHC (Myh6) gene and the gene for the regulatory non-coding RNA Myheart (Mhrt), both of which are involved in the execution of the α/β MHC switch during cardiac development and pathology. PLN^R9C^ overexpression is evident at the age of 1 month but declines dramatically along with a less pronounced concomitant decline of the resident PLN expression, until the animals die. Expression of the non-coding RNA Mhrt in PLN^R9C^ mice also exhibits a profound deregulation, despite the presence of the second, intact allele. Hence the mouse strain does not faithfully model a human PLN^R9C^ heterozygote, wherein both the mutant and the wildtype PLN alleles have, in all likelihood, the same temporal expression profile. The intricate regulatory circuit of the α/β MHC switch, involving the non-coding RNA Mhrt, was described in detail only recently, and since publications about αMHC-driven transgenes rarely contain the definition of the transgene integration site or temporal expression profile, it is suggested that some of the pathological phenomena attributed to expression of αMHC-driven transgenes may have an alternative explanation.

## Introduction

Complex genetic landscape of inherited cardiomyopathy includes many genes, among which are the genes encoding contractile proteins, such as the subunits of the myosin complex, as well as the gene encoding a small phosphoprotein phospholamban (McNally et al. 2015). Myosin is a hexameric protein complex, often called a molecular motor (Gupta 2007), consisting of two heavy chains (MHC) and four light chains (MLC). The two MHCs, designated αMHC and βMHC, are encoded by two closely spaced genes, called MYH6 and MYH7 in man, that are transcribed in the same direction (Morkin 2000). The products of individual genes combine to make three myosin isoforms V1, V2 and V3, by association of αα, αβ and ββ chains, respectively.

Expression of the two MHC gene transcripts is known to occur at a high level in all of the cardiomyocytes by 8 d p.c. in the mouse embryo (Lyons et al. 1990), when an S-shaped tubular heart has formed and contractions start (Sissman 1970). Myosin isoform switch happens more than once during development of a healthy mouse heart, and any gene put under control of the αMHC promoter should follow its expression profile in transgenic animals (Subramaniam et al. 1991). Due to its spatially restricted, cardiac-specific pattern of expression (Gulick et al. 1991), since 1991 the αMHC promoter has become the promoter of choice for transgenic expression of proteins, suspected to be involved in heart dysfunction. In hindsight, its use looked problematic from the beginning (Subramaniam et al. 1991), since only spatial, but not temporal or regulatory characteristics of the promoter were taken into consideration. However, neither the knowledge that the α/β MHC isoform switch is affected by the thyroid hormone (Gustafson et al. 1986; Izumo et al. 1986; Subramaniam et al. 1991) as well as observed in experimental cardiac hypertrophy (Morkin and Kimata 1974), nor the knowledge that the regulatory effects of hormones and experimental cardiac hypertrophy are exerted differently in ventricles and atria (Morkin 1993), had prevented the extensive use of αMHC-promoter driven transgenes in cardiovascular research (Molkentin and Robbins 2009).

Phospholamban (PLN), a 52-aminoacid transmembrane phosphoprotein, abundant in the endoplasmic reticulum of cardiac and skeletal muscle, regulates activity of sarcoendoplasmic reticulum Ca^2+^ ATPAses (SERCA). Since its discovery by Kirchberber and Katz in 1975 (Kirchberber et al. 1975) it received much attention as a possible pharmacological target in heart failure, since it is expressed in ventricles, atria (Koss et al. 1995) and sino-atrial node of the pacemaker (Maltsev et al. 2006). Transgenic expression of PLN mutants in mice has become one of the mainstream models of heart failure in the last two decades (Koss and Kranias 1996; Molkentin and Robbins 2009), reinforced by the discovery of actual, albeit rare, patients that carry PLN mutations (Fish et al. 2016; Haghighi et al. 2003; Liu et al. 2015; Schmitt et al. 2003). Recent whole genome association studies mapped the quantitative trait locus on chromosome 6q22 that includes the PLN gene, as a strong candidate for resting heart rate (Eijgelsheim et al. 2010).

Researchers in the field of PLN functional studies were among the first to embrace the transgenic technology, and a pioneering study succeeded in creating two transgenic lines, having 2 and 3 copies of the αMHC promoter–driven PLN transgene per genome, and featuring a two-fold excess of the transgenic PLN in the ventricles (Kadambi et al. 1996). These mice demonstrated no gross pathology and had normal life span.

In contrast, a mouse strain with transgenic expression of a heterozygous substitution of cysteine for arginine 9 in phospholamban (PLN^R9C^) (Schmitt et al. 2003) under the control of αMHC promoter, reported in 2003, featured dilated cardiomyopathy, heart failure and life expectancy of only 21±6 weeks. The ensuing line of physiological/genetic (Schmitt et al. 2009), transcriptomic (Burke et al. 2016) and proteomic (Gramolini et al. 2008) studies of this strain was mostly focused on Ca^2+^ dysregulation and did not address the possible impact of other mechanisms, unrelated to dysregulation of SERCA2a. Here, it is reported that the PLN^R9C^ transgenic line has a peculiar genomic structure, wherein the tandem array of 13 expression cassettes has integrated into the bi-directional promoter that drives the transcription of the resident αMHC gene (Myh6) and the gene for long non-coding RNA Mhrt (Han and Chang 2015; Han et al. 2014), involved in developmental and pathological α/β MHC switch.

## Materials and methods

### Transgenic mice and genotyping

The transgenic mouse line, carrying rabbit phospholamban with a substitution of cysteine for arginine 9 under the control of the mouse αMHC promoter (TgPLN^R9C^) was described previously (Schmitt et al. 2003). The mice were propagated by backcross to FVB/NCrl (strain code 207, Charles River Canada), thus obtaining only transgenic hemizygotes, and transgenes were identified by PCR (Zvaritch et al. 2000).

### RNA and DNA isolation

Mice were dissected under total anesthesia with diethyl ether and subsequently killed by cervical dislocation. Hearts were excised, gently squeezed while submerged in cold phosphate-buffered saline to reduce blood contamination, dissected into atria and ventricles and flash frozen in liquid nitrogen. Total RNA was extracted with Trizol reagent (Thermo Fisher Scientific # 15596-026) essentially as described by the manufacturer, subsequently treated with RNAse-free DNAse I (Thermo Fisher Scientific # AM2222) and purified by phenol extraction and ethanol precipitation. RNA from mutant and wild type litter mates (three of each genotype) were always processed in the same batch to minimize variations of quality. RNA amount and quality was estimated using Nanodrop 1000 spectrophotometer. 120 ng portions of total RNA from the mice of each genotype were pooled for cDNA synthesis. Mouse genomic DNA was isolated from either tail or ear clips as described (Miller et al. 1988) or recovered from crude preparations of heart RNA.

### Quantitative reverse transcription-assisted polymerase chain reaction

Total RNA was reverse transcribed into complementary DNA (cDNA) using Transcriptor^®^ enzyme, Roche Diagnostics, Laval, QC), primed by 10 pmol of T7-oligo-dT_24_VN primer (Thermo Fisher Scientific #AM5712) by incubation for 15 min at 42 °C followed by 1 hour at 50 °C. For the detection of long non-coding RNA the cDNA synthesis was primed with specific primers and M-MLV Reverse Transcriptase, RNase H Minus (Promega #<u>M3682</u>). A 4-µl portion of each completed cDNA reaction was diluted 1:25 with reagent-grade water and 5 µl aliquots of diluted cDNA were added to 20-µl qRT-PCR reactions, along with 5 µl of a 1 µM primer pair and 10 µl of SYBR^®^ Select reagent master mix (Applied Biosystems #4472919). A “no primer” control reaction was analyzed in parallel with properly made cDNAs. The identity of all fragments used in real-time PCR was verified by sequencing.

Reactions were run in triplicate on the Mini-Opticon qRT-PCR instrument (Bio-Rad Laboratories Canada Ltd.). The hypoxanthine phosphoribosyl transferase gene was used as a reference. The qRT-PCR Ct values were exported as Excel files and analyzed using online software provided by Qiagen Inc [http://pcrdataanalysis.sabiosciences.com], which included two-tailed Student’s t-test, and finally converted into a graph using GraphPad Prism v.6. Data were considered significant with p-value less than 0.05.

### Determination of the ratio of native to transgenic PLN transcripts

PLN- and TgPLN^R9C^ primer pairs were used to amplify the respective PCR fragments from cDNA by conventional PCR, which were purified and quantitated. After two million-fold dilution of each fragment in a glycogen solution (20 µg/ml), synthetic molar ratios of 5:1, 2:1, 1:1, 1:2 and 1:5 were made and each mixture was tested with the same two non-overlapping primer pairs in a quantitative PCR reaction. A calibration curve, plotting the function of constructed ratio to experimentally observed ratio was thus obtained which was used to correct the experimentally observed ratio in a tissue. Thus, the ratio of 2^−ΔΔCt^ of PLN relative to TgPLN^R9C^ can be adjusted for primer efficiency according to the formula y=4.4136x-0.0855, where *x* is the experimentally observed ratio, and *y* is the corrected ratio. By design, this formula applies only to the two specific primer batches that were actually used, and the procedure needs to be repeated for any new primer batches.

### DNA amplification and sequence analysis

Genomic location of the transgenic insert was determined by inverse PCR (Does et al. 1991). Genomic DNA (0.5 µg) was cut with Nco I (one of the sites was present at the start codon of the TgPLN^R9C^ gene but not elsewhere within its transcript), diluted to 700 µl and circularized with T4 DNA ligase overnight at room temperature. The DNA was phenol extracted and ethanol precipitated and dissolved in reagent grade water at a concentration of 25 ng/µl. Primers P1 and P2, specific for the rabbit TgPLN^R9C^ gene, directed outward, were used to amplify the circularized fragment. The location of the junction sequence, deduced from sequencing of the fragment, was verified by regular PCR using appropriate flanking primers P1-P4 and P3-P5 (Fig. 3a and 4). Rapid amplification of cDNA 3’-ends was done with T7-oligo-dT_24_V primed cDNA, using PCR with anchor-specific primer and gene-specific primer P1. Due to high concentration of the PLN transcript it was readily amplified by a single round PCR. If purified genomic DNA was not available, genomic DNA amplification could be successfully carried out from the crude heart RNA preparations (not treated with DNAse). Amplifications were carried out either with Platinum^®^ Taq polymerase (Thermofisher) or with LongAmp^®^ Taq polymerase (New England Biolabs) on a Bio-Rad C1000 Touch™ thermal cycler. All PCR products were analyzed by agarose gel electrophoresis, purified on Qiaquick MinElute columns (Qiagen) and sequenced at the Toronto Centre for Applied Genomics facility. Sequence assembly was done with Sequencher 4.10 (Gene Codes, Ann Arbor, MI). Reference *Mus musculus* C57BL/6J genomic and cDNA sequences were retrieved from Ensembl [http://www.ensembl.org]. Oligonucleotide primers were synthetized by ACGT (Toronto, ON, Canada) or Eurofins MWG Operon LLC (Huntsville, AL, USA). Primer sequences are listed in Table 1.

**Table 1.**
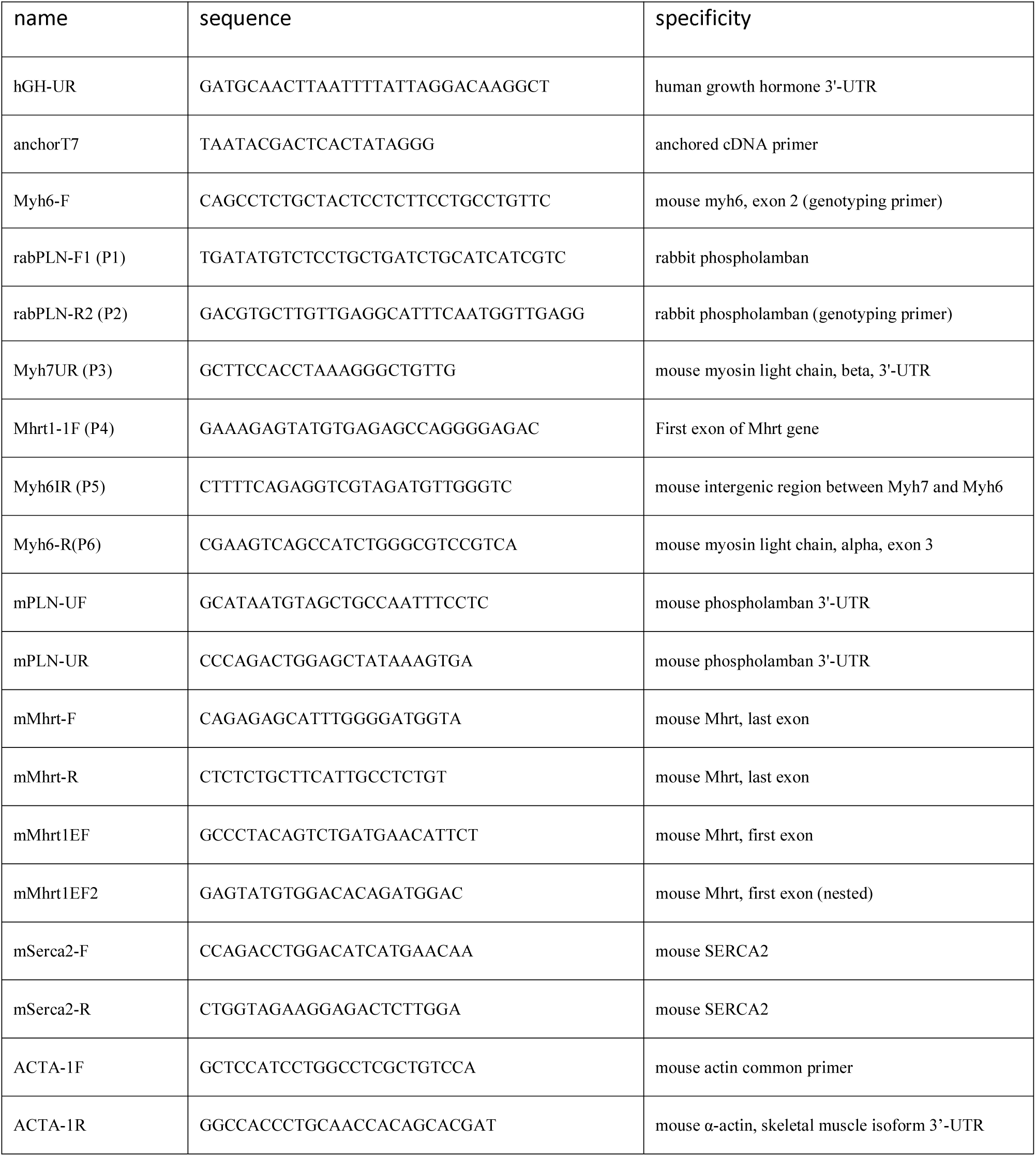
Synthetic oligonucleotides used in the study

## Results

### Developmental suppression of PLN^R9C^ expression in transgenic hearts

Overexpression of wildtype PLN, if driven by the αMHC promoter, is expected to disturb the natural atrio-ventricular PLN gradient (Koss et al. 1995), wherein 3 to 5 fold more PLN is found in normal murine ventricles than in atria; besides, the expression of αMHC begins somewhat earlier in atria than in ventricles (Lyons et al. 1990). In a wildtype mouse the activity of the αMHC promoter should peak between 1 and 2 months of age (Han et al. 2014; Ng et al. 1991), while βMHC expression drops to a very low level after birth (Ng et al. 1991). Therefore, it was first sought to compare gene expression of the transgenic and native PLN at 1 month. Using real-time PCR, the ratio of TgPLN^R9C^ to native PLN transcripts at 1 month was found to be slightly above 6-fold (Fig.1a), while native PLN in transgenic hearts was found downregulated 1.8-fold relative to the wildtype (Fig.1d). If the level of TgPLN^R9C^ overexpression at 1 month is maintained in translated PLN protein, a ratio required to trap the native PLN in a non-productive pentameric complex, where it cannot be phosphorylated by protein kinase A (Ha et al. 2011; Wittmann et al. 2015) is substantially exceeded.

**Fig. 1.**
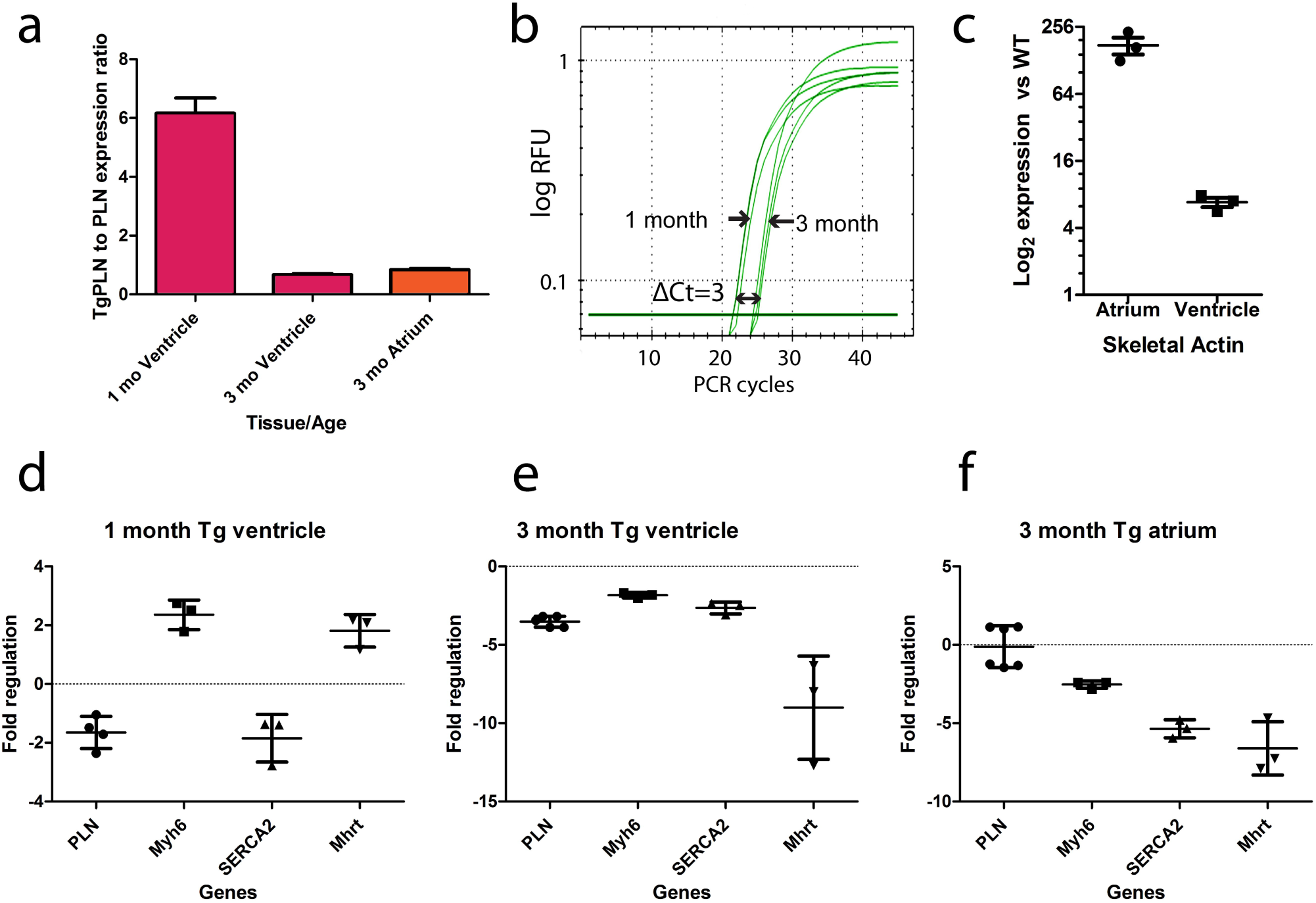
Genotype-specific and developmental changes in gene expression in the hearts of transgenic mice expressing mutant PLN. **a**: Expression ratios of transgenic PLN to resident PLN in transgenic hearts only **b**: Quantitative PCR curves, showing dramatic downregulation of transgenic PLN in postnatal development of transgenic ventricle **c**: Upregulation of skeletal actin in 3-month old transgenic heart relative to wildtype heart **d-f**: Expression levels of selected genes in transgenic heart relative to wildtype heart. P<0.05 for all inter-genotype comparisons

TgPLN^R9C^ animals start to die (Gramolini et al. 2008; Schmitt et al. 2009), at about 3 months of age. Gene expression experiments done with the same starting amount of RNA, allowing a direct comparison of Ct values, show that absolute expression of TgPLN^R9C^ goes down ~8 fold in the timeline from 1 month to 3 months (Fig. 1b). The ratio of TgPLN^R9C^ to native PLN transcript in the 3 month ventricle is only 0.69 (Fig.1a). Evidently, at 3 months there is no overexpression of TgPLN^R9C^ in the ventricle anymore, similarly to previous reports (Babu et al. 2006; James et al. 2000; Kadambi et al. 1996; Mende et al. 1998; Nakayama et al. 2003; Sanbe et al. 1999; Zvaritch et al. 2000), even though the native PLN is downregulated 3.7 times relative to the wildtype (Fig.1e). Presumably, somewhere at an intermediate age between 1 and 3 months the transgenic PLN expression transiently passes the 1:1 ratio with the native PLN, to model a situation thought to exist continuously in a human patient heterozygous for the PLN mutation. In contrast, the TgPLN^R9C^ heart initially receives a large excess of mutant PLN around the age of 1 month, but the transgene expression then gradually subsides on the background of developing cardiomyopathy, until the animal dies. The documented use of a similar construct to create transgenic mice may result in dramatic suppression (Mende et al. 1998), as well as in no suppression (Santini et al. 2007) of the transgene. A peculiar feature of the 3-month TgPLN^R9C^ heart is a strong upregulation of skeletal actin, especially in the atrium (over 160-fold, Fig.1c), incidentally, also observed in heterozygous mice with αMHC gene ablation (Jones et al. 1996), suggests that overexpression of TgPLN^R9C^ may not be the only factor of the morbid phenotype.

### Determination of the chromosomal structure and location of the transgene

In search for the reasons of peculiar changes in the gene expression of the TgPLN^R9C^ mouse heart, in particular of the dramatic downregulation of the TgPLN^R9C^, the genomic sequence of the TgPLN^R9C^ chimeric gene and also the sequence of its transcript were determined from appropriate PCR products but no potentially detrimental structural alterations were found. By using primers P1 and P2 (Fig.3a), specific to the TgPLN^R9C^ gene and facing outward in a long-range PCR, it was confirmed that each unit contains the expected 5.5 kb fragment containing the αMHC promoter [Genbank Accession U71441] and exists exclusively in a head-to-tail orientation in a tandem array, since a single product (Fig.2a, leftmost panel) with an unambiguous sequence reading across the unit junction was obtained [Genbank Accession KU665646]. Using real-time PCR on genomic DNA with αMHC (Myh6) gene as reference, it was also found in one randomly chosen animal, using Myh6 gene as a reference, that the transgenic array contains 13 copies of the expression cassette (not shown).

**Fig. 2.**
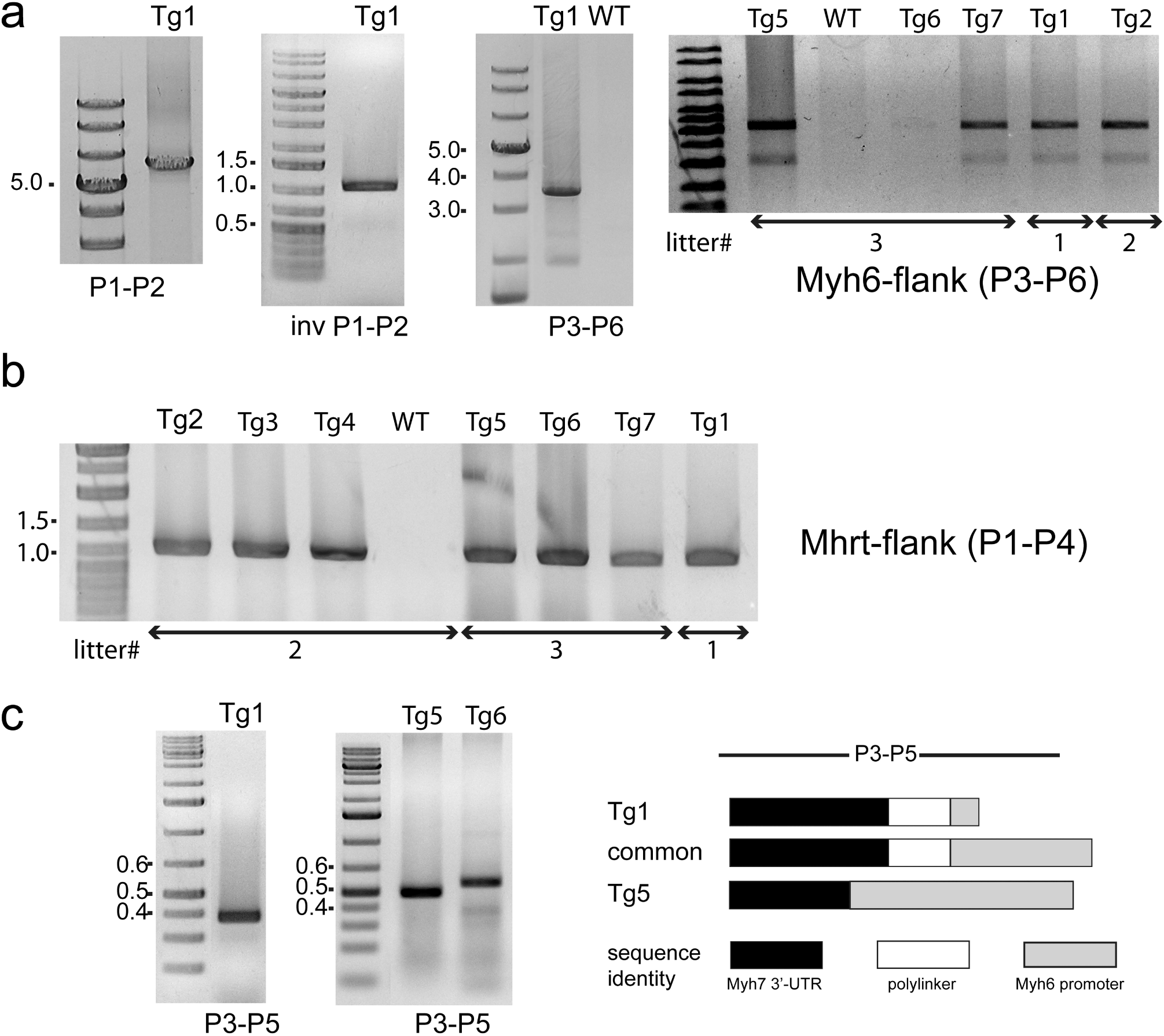
Results of PCR experiments used to determine the structure and the chromosomal location of the transgenic array. **a**: From left to right, PCR amplification with outwardly directed primers across two adjacent repetitive units; inverse PCR amplification of genomic DNA treated with Nco I and ligase; long range PCR amplification defining the distance of the transgenic array from Myh6 exon 3 **b**: PCR amplification defining the distance of the transgenic array from the Mhrt exon 1. **c**: Artefactual deletion of junctional sequences in short PCR products spanning the transgenic/genomic junction. Numbers beside the gel panels denote the marker fragment length in kilobase pairs, Tg1 to Tg7 denote transgenic animals, and P1 to P6 denote PCR primers, listed in Table 1. Litters are numbered in chronological order, starting from #1, but are not adjacent generations

**Fig. 3.**
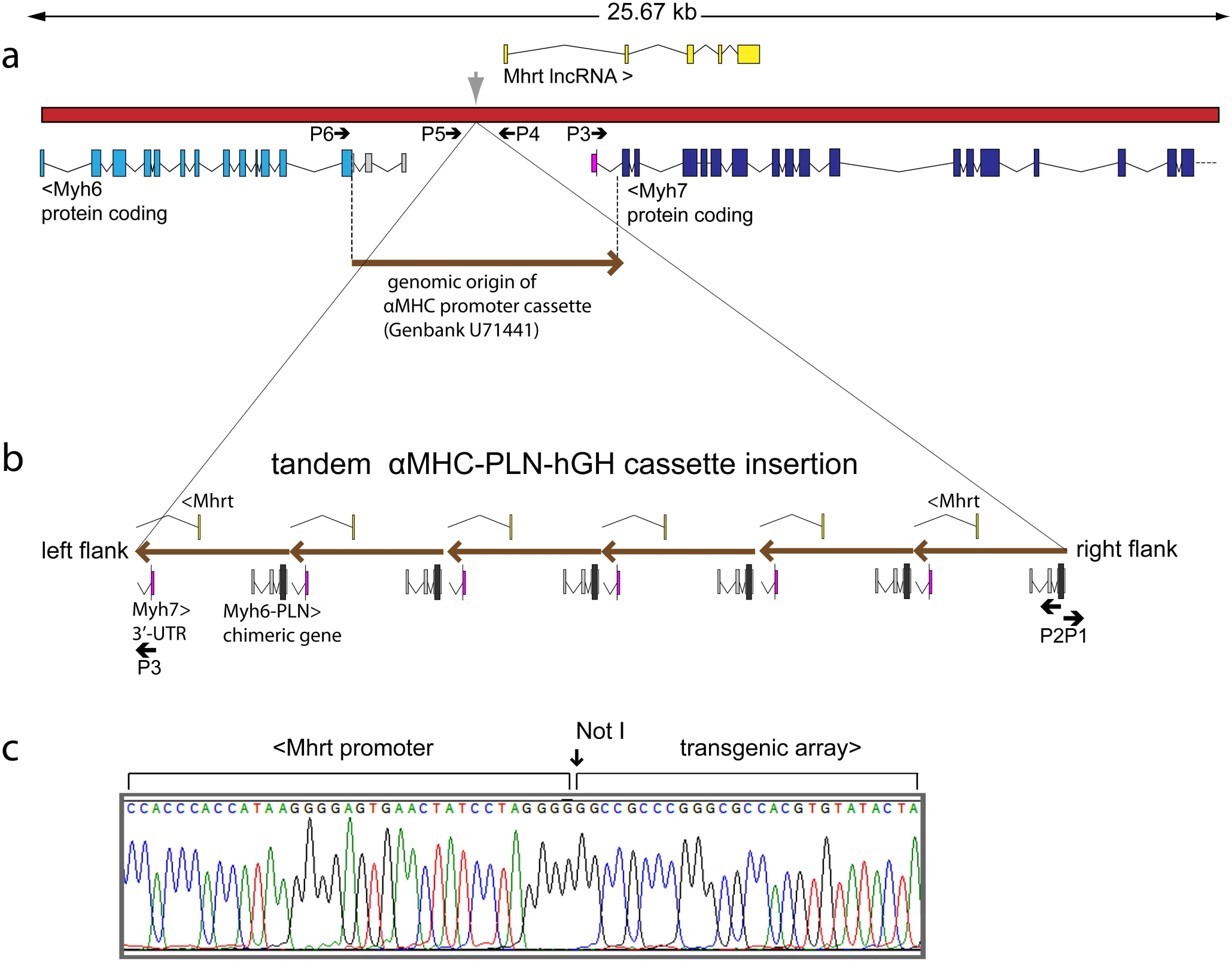
A portion of mouse genome on chromosome 14, showing the origin of αMHC promoter fragment and the transgenic array integration point (*grey arrow*), as deduced from PCR and sequencing experiments on genomic DNA. **a**: Relative locations and direction of transcription of the genes encoding α-myosin heavy chain (Myh6), β-myosin heavy chain (Myh7) and long non-coding RNA Myheart (Mhrt). **b**: Deduced structure of the transgenic array. One complete expression cassette is shown as a *long arrow*. Fewer cassettes are shown for clarity. Short arrows denote the location of oligonucleotide primers P1 to P6, listed in Table 1. **c**: Junctional sequence of the transgenic array and the mouse genomic DNA, facing the Mhrt gene, from a PCR fragment generated with primers P1 and P4 (Fig.3 and Table 1)

**Fig. 4.**
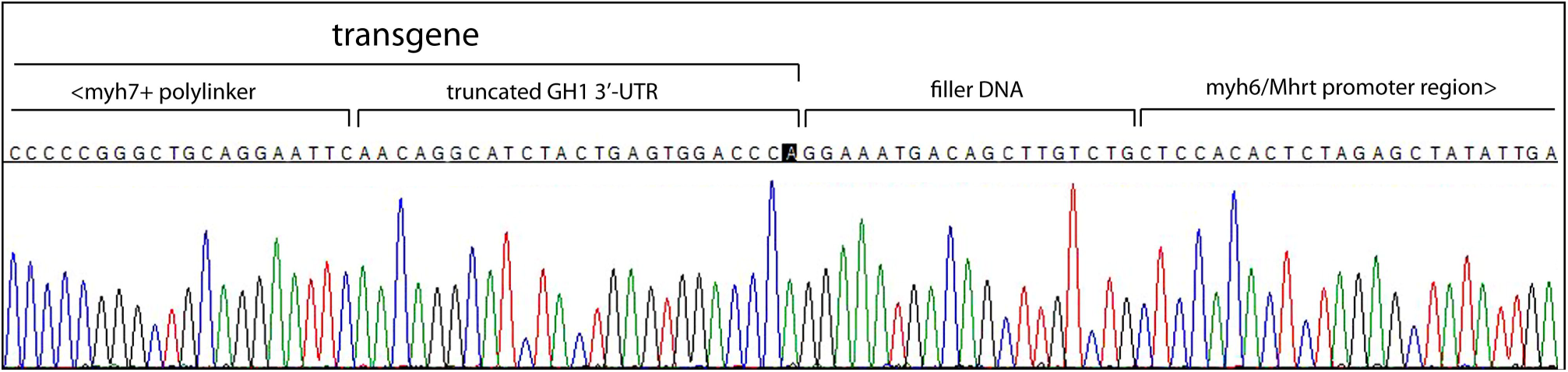
Junctional sequence of the transgenic array and the mouse genomic DNA, facing the Myh6 gene, from a PCR fragment generated with primers P3 and P5 (Fig. 3 and Table 1)

Chromosomal location was determined with appropriate primers in the conventional and the inverse PCR format (Fig.2a). Determination of the side of the transgenic array, facing the Mhrt gene, presented no technical problems and it was found to be identical in 7 transgenic animals (3,3,1) from three different litters (genomic coordinate 14:54968214 of Mouse Genome assembly GRCm38.p4 (GCA_000001635.6)). The transgene portion contains a filled-in Not I site of a polylinker sequence (Fig.2b ad Fig.3c), presumably used to excise the construct from the vector (Schmitt et al. 2003), indicating that it is the original end of the construct (Pawlik et al. 1995). Determination of the flanking sequence at the other end of the transgene presented certain technical problems. Similarly, 7 transgenic animals from three different litters were analyzed. Initially, a flanking primer (P5) was designed to be close to the projected junctional sequence from the results of the inverse PCR and paired with a primer specific for the Myh7 3’- UTR in the transgene (P3). The first use of this primer pair recovered products only in mice Tg1 (one litter) and Tg5, Tg6 (another litter), which were of slightly different size (Fig.2c). Absence of product in other animals could be due to a deletion affecting one of the primer sites or due to a PCR failure. However, by using other primers and conditions junctional sequences were eventually obtained from the remaining animals. The common junctional sequence contains a double polylinker, identical to the one in KU665646 and 26 bp of the sequence derived from the human growth hormone gene. Thus, it corresponds to a severely truncated original cassette (Fig. 4). The truncated transgenic unit, however, is not directly juxtaposed to mouse genomic DNA, corresponding to the aMHC promoter, but is separated from it by a 19-bp “filler” sequence (Mayerhofer et al. 1991), the origin of which could not be established. Comparison of junctional sequences revealed that some of them contain short internal deletions (Fig.2c) relative to a common sequence, as originally found in animals Tg1 and Tg5. Finally, long range PCR with the primer P3 paired with a primer placed on Myh6 exon 3 outside of the portion included in the transgene (P6) produced a fragment of expected size from all litters (Fig. 2a) that is consistent with the location of the junctional sequence facing the Mhrt gene. An attempt to sequence the long range PCR product across the junction encountered a sequence heterogeneity (not shown) around the location of the polylinker and supported the conclusion that the small deletions, observed in short PCR products, are likely PCR artefacts, induced by a local secondary structure. Thus, no extensive rearrangement or large genomic deletions accompanying the transgenic insertion were found, unlike those reported for other transgenes (1, 2).

Taken together, these experiments showed that the transgenic array is integrated into the bi-directional promoter (Haddad et al. 2003; Han et al. 2014) driving the resident Myh6 gene and the resident long non-coding RNA Mhrt in such a way that the TgPLN^R9C^ transcription is driven by the transgenic copies of the promoter in the opposite direction to that of the resident Myh6 gene (Fig.3b). The insertion of the transgene results in 83 bp deletion of the target site in the mice analyzed. The resident Myh6/Mhrt promoter appears to be cut approximately in half by the ~85 kb transgenic array insertion, while the other Myh6 allele is intact, judging from the ongoing transcription of Myh6, verified by real-time PCR (Fig.1d-1f). From the small number of animals available for analysis no definite conclusion about the transgene stability could be made.

### Deregulation of non-coding RNA Mhrt expression in 3 month old transgenic hearts reflects insertional mutagenesis of its promoter

Myh6 promoter substantially overlaps with the promoter of the long non-coding RNA Mhrt (Han et al. 2014), thus it was of interest to determine if the transgenic insertion had indeed any influence on its expression. Using a pair of primers, specific for the 3’-end of Mhrt in a quantitative PCR format, about 2-fold upregulation was observed in transgenic ventricle at 1 month and over five-fold downregulation both in transgenic ventricles and atria at 3 months (Fig.1d-1f), wherein the level of Mhrt expression drops close to a reliable limit of detection. A confirmatory end-to-end amplification and sequencing of Mhrt was done from both transgenic and wildtype ventricle cDNA, using one of the 3’-end specific primers utilized for cDNA synthesis and quantitative PCR and a nested pair of primers, specific for the first exon of Mhrt (Table 1).

The deregulation of Mhrt expression occurs despite the presence of the intact allele in the hemizygote, as if the damaged allele exerts a dominant negative effect over the intact allele.

## Discussion

The peculiar genomic structure of the TgPLN^R9C^ mouse line, containing damaged Myh6/Mhrt allele and presumably impaired α/β MHC switch is a probable contributing cause of the morbid phenotype of this line, along with the excessive amount of mutant PLN that the mouse heart receives during the 1st month after birth. Indeed, only a weak downregulation of αMHC (this study) and upregulation of βMHC (Schmitt et al. 2009) is observed, and the non-coding RNA Mhrt is paradoxically downregulated on the background of the developing cardiomyopathy (Han et al. 2014). Even in the absence of data about gene expression in humans, it can be concluded that this line does not faithfully model a heterozygous human patient carrying a cysteine for arginine 9 mutation in PLN, as the latter not only expresses the mutant and the wildtype PLN from identical promoters, but also does not carry a Myh6/Mhrt allele damaged by insertional mutagenesis. While it may be argued that the transgene localization, as presented here, is but an elaborate PCR artefact, however, it is supported by the observed deregulation of Mhrt that is a predicted consequence of the damage to its promoter.

It can also be argued that the transgene structure as revealed in this study may not necessarily have existed in the founder transgenic mouse, but may have appeared as a result of a secondary transgene rearrangement (or several ones) at some point in 12 years of conventional breeding. It has become customary to cite Aigner and colleagues (Aigner et al. 1999) who did a thorough study to prove that transgenes, once obtained, are stable over many generations. However, their conclusions from a study of just one gene appear to be an unjustified generalization and may not apply to all conceivable transgenes. Specifically, there are two important differences between their case and that of TgPLN^R9C^. First, TgPLN^R9C^ animals have severely reduced viability; hence a spontaneous transgenic array expansion (Barlow et al. 1995), contraction (Alexander et al. 2004) or even change of location (Kearns et al. 2001) may be selected for if it leads to a weaker morbid phenotype. Second, the transgenic array is inserted in an inverted orientation into a homologous sequence, which makes it prone to instability. Furthermore, the current genotyping protocol would not reveal most of the interunit rearrangements, while only a complete transgene excision will be genotyped as wildtype and discarded. However, in the absence of extensive structural data in a large pedigree no definite conclusion can be made. The issue of the PLN^R9C^ transgene stability may be an interesting topic for a future dedicated study.

In the past, the problem of transgene silencing was well recognized (Garrick et al. 1998; Martin and Whitelaw 1996; Robertson et al. 1995; Sanbe et al. 2003; Sutherland et al. 2000), so obtaining a clinically relevant phenotype in a TgPLN^R9C^ mouse line could have been a reason for its initial selection. However, even though the case of the TgPLN^R9C^ mouse line may be rare or even unique, it brings to light two issues of general importance for transgenic research that, in this author’s opinion, supersede the importance of mechanistic insights derived from the morbid phenotype of a mouse line allegedly modelling a rare cause of cardiomyopathy. One is the lingering lack of attention to the transgene integration site, an apparent relic of the “pre-genomic era”, quite surprising in view of the current availability of the refined mouse genomic sequence (Church et al. 2009). The other is the far-reaching implications of the recently defined regulatory elements of the α/β MHC switch (Han et al. 2014; Hang et al. 2010) to the interpretation of data collected over 25 years of use of αMHC-driven transgenes.

Determination of transgene integration site is inherent to the gene trap methodology (Friedrich and Soriano 1991), which in essence is an ingenuous exploitation of a drawback of unguided transgenesis (Wigler et al. 1977). However, it is disconcerting to find that the majority of cardiovascular researchers working with mice obtained by unguided transgenesis are apparently not concerned with the issue of transgene integration site. Even though it clearly does not cover all relevant studies, a search in PubMed database for titles containing ”cardiac-specific overexpression” found 81 hits from 1997 to 2015 (as of April 2016), wherein 72 studies used the αMHC promoter (Subramaniam et al. 1991). Further search with “*cardiomyocyte*-specific overexpression” found additional 17 references starting from 2012, of which 8 used αMHC promoter and 8 used binary/inducible systems (Sanbe et al. 2003; Sohal et al. 2001; Yu et al. 1996). However, only one study of the 98 used both targeting to a specific locus and inducible expression with a binary system (Schuster et al. 2016) and *none* actually determined the transgene integration site, regardless of whether the mouse line was novel or re-used, while the mouse phenotype as a model of a human disease was always addressed in detail (see Supplementary References).

Some of the pitfalls of the process of transgene creation have been known for quite a long time (Bronson and Smithies 1994; Robertson et al. 1995) and were difficult to avoid in the absence of the complete genomic sequence. In 1992 Meisler stated that about 5% of transgenic insertions damaged a gene (Meisler 1992). Detrimental consequences of an unguided insertion occasionally lead to discoveries of new genes (Krakowsky et al. 1993; Maguire et al. 2014). However, after nearly 40 years the appreciation of chances of transgene causing complex disruption of a genomic region (West et al. 2016) has grown considerably, since now we are well aware of the ubiquitous presence of non-coding RNA genes, intricately entangled with protein-coding genes (Mattick and Makunin 2006), as well as of long-range enhancers (Smallwood and Ren 2013).

Furthermore, the presence of a transgene should also pass the test of non-interference with embryonic development. Here, the αMHC promoter stands apart from many other promoters utilized in transgenic research in that, unbeknown to its users for decades, it contains a truncated regulatory circuit, involving a long non-coding RNA, Myheart (Haddad et al. 2003; Han et al. 2014). Revisiting the pioneering study of the αMHC promoter as a driver in transgenic mice (Subramaniam et al. 1991), one finds an interesting fact: of the 17 lines generated, 9 were from the 5.5 kb construct (the αMHC promoter), 7 from the 1.3 kb construct (reportedly inactive), and one from the 3 kb construct. Curiously, the αMHC gene-distal border of the 3 kb construct (Sph I site at nucleotide 2656 of Genbank Accession U71441) is very close to the site of the 85-kb insertion (at nucleotide 2732 of U71441), observed in the TgPLN^R9C^ mouse line. Thus, in the latter, one Myh6 allele is effectively driven by the equivalent of the 3 kb construct (Subramaniam et al. 1991), with the adjacent transgenic sequence containing an inverted repeat of it. It is probable that any mice displaying a morbid phenotype in the context of the study of a protein having no known function in mouse (chloramphenicol acetyl transferase), would have been discarded. To the best of this author’s knowledge, the 3 kb αMHC promoter construct was never used again in stably transformed transgenic mice.

25 years of successful outcomes of transgenic expression of *functionally relevant* proteins under the αMHC promoter feature a wide spectrum of examples: from just one breeder line, a minimal requirement for publication, with one or a few copies of transgene per genome and an unimpressive level of overexpression, up to generation of multiple lines with a wide range of the number of copies per genome and/or of transgene expression rate, sometimes far exceeding that of the resident gene (examples are found in the Supplementary References).

Even though the parameters of transgenic strains, such as transgene integration site, copy number or expression rate may span a wide range of values (where determined), the variety should not necessarily prompt a conclusion that the parameters are independent of each other or of the transgene function, and are determined only by chance. If chance were the main factor, the use of the original αMHC promoter was remarkably successful in view of the possible insertional mutagenesis, as only a few disconcerting events have been documented. For example, it was found that green fluorescent protein gene expression driven by the αMHC promoter could cause dilated cardiomyopathy (Huang et al. 2000) and that the αMHC promoter driven Cre recombinase (Sohal et al. 2001) should be carefully controlled (Lexow et al. 2013) to avoid cardiac fibrosis (Koitabashi et al. 2009). Theoretically, once such facts have come to light, every author should be defending his/her position that any αMHC-driven transgenic phenotype is not an artifact, i.e. by presenting appropriate controls (Huang et al. 2000). Instead, a mere possibility of an artifact is confidently dismissed. It took 15 years to reveal that the αMHC-driven Cre recombinase transgene (Sohal et al. 2001, prone to induction of cardiac fibrosis, has, in fact, produced a dramatic genomic rearrangement {Harkins, 2016 #254).

10 years after the αMHC promoter introduction, Krenz and Robbins, commenting on the results of transgenic studies of the Bcl-2 gene in an article entitled “Gates of Fate” (Krenz and Robbins 2001) admitted that one “cannot differentiate between the possibilities of either direct causality, or whether we are simply observing a secondary, compensatory effect due to transgenically mediated overexpression”. Incidentally, the same applies to the TgPLN^R9C^ mouse line. While the allusion to fate points to the role of the Bcl-2 protein in cell death, from today’s perspective the αMHC promoter itself may have more to do with the Gates of Fate than currently appreciated, as it is a site of remarkable phenomena that take place during embryogenesis and after birth, by virtue of containing a binding site for a nucleosome remodeling complex with DNA-helicase Brg1 (Khavari et al. 1993; Randazzo et al. 1994) and an overlap with the promoter for regulatory long non-coding RNA Mhrt (Han et al. 2014).

As mentioned earlier, in wildtype mouse the αMHC transcript achieves maximal postnatal level between 1 and 2 months after birth (Ng et al. 1991) and then declines, however, one does not often find studies of the temporal αMHC-driven transgene expression profile (Mende et al. 1998; Santini et al. 2007; Sheridan et al. 2000). The relatively few studies that did it, have uncovered that transgene expression falls into two principal scenarios: one where it closely follows the expression profile of the αMHC promoter in the wildtype mouse (Santini et al. 2007), and another where its rate first increases and then drops to a level close or even below that of the wildtype at the same age ((Mende et al. 1998) and this study). In some cases, the transgene suppression is not evident, but appears after the mechanical stress is applied to the heart (Sheridan et al. 2000). Many studies, where the overexpression level was measured only once at a relatively advanced age, report a value not exceeding 2-fold excess over the resident protein level (Babu et al. 2006; James et al. 2000; Kadambi et al. 1996; Nakayama et al. 2003; Sanbe et al. 1999; Zvaritch et al. 2000).

The transcriptional suppression can be explained by the fact that Brg1 is normally expressed in embryonic but not in adult cardiomyocytes (Hang et al. 2010). Pro-hypertrophic stimuli can induce its expression, resulting in an attempt to execute a pathological α>β MHC isoform switch (Papait and Condorelli 2010). Any αMHC transgenic array has multiple binding sites for Brg1, thus it may enforce a quantitative (partial) failure of α>β switch under stress conditions and eventually cardiac failure on a macroscopic level, e.g. due to uncoordinated remodeling of ventricles and atria. The cardiac failure would be promptly attributed to the action of the transgene (such as TgPLN^R9C^), while actually it could be a combination of at least two factors and in some cases could have nothing to do with the transgenically expressed protein at all. As noted in (Huang et al. 2000), the transgenes that behave according to this scenario are actually presented without *appropriate* controls (which may amount to much additional work), thus observations based on such studies may do little beside skewing the knowledge space related to cardiac pathology.

The native αMHC promoter is one of the targets of Brg1-containing nucleosome-remodeling SWI/SNF complex (Whitehouse et al. 1999) throughout the development and in the postnatal life (Hang et al. 2010), and it is conceivable that its supernumerary copies in a transgenic array may interfere with the availability of Brg1. Hence, there appears to be an entirely novel aspect of the αMHC promoter in a transgenic environment, albeit one of a more speculative nature. Mammalian SWI/SNF complexes utilize one of the two highly similar ATP-dependent DNA helicases Brm (Brahma, also known as SMARCA2) or Brg1 (Brahma-related gene 1, also known as SMARCA4) as the catalytic subunit (de la Serna et al. 2006; Ho and Crabtree 2010). Upon association with promoters by sequence-specific transcription factors, SWI/SNF complexes displace nucleosomes away from transcriptional start sites to allow RNA Polymerase II access and initiation of transcription. Cardiogenic transcription factors TBX5, GATA4, and Nkx2–5 interact with SWI/SNF to program non-cardiac mesoderm into cardiomyocytes (Bruneau 2010). SWI/SNF complexes participate in many diverse developmental processes (reviewed in (Smith-Roe and Bultman 2013)).

Brg1 is expressed at much higher levels than Brm in the early embryo (LeGouy et al. 1998), and the embryonic stem cell SWI/SNF complex (esBAF) incorporates Brg1 as the catalytic subunit (Ho et al. 2009). However, Brm participates in a subset of developmental processes that are sensitive to Brg1 dosage; when the combined gene dosage is diminished from four copies in wild-type (Brg1+/+;Brm+/+) embryos to one copy in Brg1+/-;Brm-/-double mutant embryos, it drops below a threshold required for implantation to occur (Smith-Roe and Bultman 2013). Conceivably, fluctuations in Brg1 availability may alter many developmental processes.

Long non-coding RNA Mhrt was known to be involved in the execution of the α/β myosin switch for over a decade (Haddad et al. 2003), but only recently a detailed study has defined its regulatory mode in chromatin remodeling (Han et al. 2014). According to it, Mhrt competes with Brg1 in regulation of transcription of both αMHC gene and its own gene from the bidirectional promoter (Han et al. 2014). Multiple copies of the αMHC promoter in a transgene are equipped with intact Mhrt promoter, but not full-length Mhrt gene itself, so the truncated copies may interfere with the execution of the α/β regulatory switch. Hence, the Mhrt/Brg1 circuit may actually contribute to the fate of αMHC transgenes, i.e. determine at multiple points in time which of them are embryonic lethal; in other words, determine which of the genes may ultimately be considered essential for heart development and pathology from the results of their transgenic expression. For example, from day 8 pc the starting position of the α/β myosin heavy chain switch is set to favor the expression of βMHC over αMHC, while after birth the toggle quickly reverses. Brg1 and Mhrt play active (and opposing) roles in each case (Hang et al. 2010). After birth the surviving transgene-carrying embryos are subject to a different kind of selection, when expression of the transgene, driven by the αMHC promoter, may reach a level far greater (Ng et al. 1991), than that of the respective resident gene (a true overexpression), with downstream effects that may have little to do with its original function. Admittedly, similar arguments have already spurred the development of more sophisticated transgenic systems in the early 21^st^ century (discussed in (Sanbe et al. 2003; Tasic et al. 2011)).

In conclusion, the complex chromatin-remodeling machinery associated with the αMHC promoter implies that in its use not only the transgene integration site and the number of copies in the unguided transgenesis are likely not due to chance alone, moreover, the effects in adult animals may only demonstrate how they cope with a protein, fluctuating excess of which is not under investigator’s control. It could be reminded that the αMHC promoter was originally selected not only for its spatially restricted activity, but also for the specific goals of the Robbins laboratory that required high level transgenic expression (Robbins 1997). Subsequently, however, it was used to express proteins with an inherently low and/or tightly controlled level of expression. Regrettably, there is no information about genes that failed to be expressed by unguided transgenesis with this promoter.

Currently, it is widely accepted (a recent review, (Nandi and Mishra 2015)) that fetal reprogramming is one of the common features of the failing heart. Re-expression of one or more of the genes normally active in the fetus is often a part of αMHC-driven transgenic phenotype, however, in view of the arguments presented above, the similarity to cardiac disease or to data from non-transgenic models should be taken with a grain of salt. Now, with the new data about the functional elements of the αMHC promoter, the insights obtained from some of such transgenes may be in need of reconsideration. In particular, the existing αMHC transgenes need to be further characterized with experiments outlined here, i.e. with regard to transgene insertion point, *temporal* transgene expression profile and expression of Mhrt. Furthermore, any gross discrepancies between the phenotype of human patient and its mouse model, such as lethality in one case and its absence in the other, should not be taken lightly and dismissed as a species difference. A successful transgene may have passed through the Gates of Fate, but if the end result represents a faithful model of the disease under study may still be a matter of debate. Fortunately, today an impressive palette of alternative genome modification strategies is available to verify these data (Giraldo and Montoliu 2001; Sanbe et al. 2003; Van Keuren et al. 2009; Woltjen et al. 2011; Yu et al. 1996). In addition, while mouse may serve well as an interim model, genome targeting of another animal may be considered as an alternative, using the CRISPR/Cas9 technology (Seruggia and Montoliu 2014).

## Conclusions and perspectives

Methods of transgene creation have come a long way since the first pioneering experiments of the 1970s, and so did the techniques of determining chromosomal localization and profiling of gene expression. A fundamentally new, no longer protein-centric, picture of the mammalian genome is slowly emerging, to which transgenic studies have made a crucial contribution. It would be unfair to pass judgement on the reasoning behind the strategy to create the PLN^R9C^ transgene that was devised over 15 years ago, in particular, to remark that the goal of modeling a human heterozygote could have been better served with a conventional knock-in. Nevertheless, it is plausible that the entire field of PLN functional research may have taken a different route, were the details of the genomic location of rather numerous PLN transgenes addressed early in their respective projects. Today not only the techniques of transgene characterization (review, (Stefano et al. 2016)) have become substantially more accessible and affordable, but also the results of genomic mapping can be compared to a refined genome sequence and its functional annotation. They may immediately alert the researcher of any possible adverse genomic effects before physiological or behavioral studies may begin. This recommendation is almost 20 years old (Robbins 1997), and there is even more incentive to follow it today.

Making the determination of the transgene parameters such as the genomic location and copy number mandatory for publication would bring unguided transgenes on par with the high standards used in the current generations of genetically modified mice, produced by large scale projects (Ryder et al. 2013). It would also be extremely helpful to routinely obtain at least a crude *temporal* profile of transgene expression. The presence of an aberrant temporal profile would then alert researcher to the possibility of artefactual phenomena on the physiological level.

The notion that the popular αMHC promoter in its original form spells trouble for studies of experimentally induced cardiac pathology is less evident, since the importance of long non-coding RNAs only recently began to gain recognition in cardiovascular research (De Windt and Thum 2015). The fact that one of these RNAs is a component of the αMHC promoter construct and hence is included, possibly in multiple copies, in many existing transgenes, means that reevaluation of the transgenes may be necessary, especially in those cases where the transgene features a cardiac pathology. Advanced approaches, such as creating a transgene with the native promoter on a large genomic fragment (Van Keuren et al. 2009), targeting transgene to a specific location (Tasic et al. 2011) and reversible transgene insertion (Woltjen et al. 2011) are available to verify the transgenic phenotype. In cardiovascular research, there is clearly a need for development of alternative tissue-specific promoters that could be used instead of the αMHC promoter in transgene creation. The concept of targeting a transgene to a pre-defined location, a safe “landing pad” (Soriano 1999; Zhu et al. 2014), can be expanded to locations where an insertion has minimal off-target effects, utilizing the data from genomic sequence and large-scale gene-trap projects (West et al. 2016; West et al. 2015).

From a more general perspective, the case of the TgPLN^R9C^, as well as of the αMHC promoter, discussed here, are examples of a downside of shared resources. Availability of diverse shared resources, such as genetically modified mice or gene expression cassettes, is an undisputable advantage of mouse research as compared to research in other laboratory animals. However, hidden faults of the shared genetics resources are similar to the bugs of widely used computer software, in that they may insidiously affect a large number of users. Therefore, the importance of data provenance in biomedical research, transgenic or otherwise, should not be underrated.

## Acknowledgements

The Author is grateful to Dr. Elena Zvaritch for her generous help with mouse surgery and for enlightening discussions. This work did not receive a dedicated funding, but it used resources from another project in which the Author participated under the direction of Dr. David H. MacLennan, whose *de facto* financial support as well as valuable comments are gratefully acknowledged. University of Toronto is acknowledged for online library access, used in the manuscript preparation, however, the views expressed in this article are those of the Author, and do not necessarily reflect endorsement by University of Toronto, or by any of its constituent departments and units.

## Ethical Approval

All applicable international, national, and/or institutional guidelines for the care and use of animals were followed. All procedures performed in studies involving animals were in accordance with the ethical standards of the University of Toronto.

